# Natural selection acting on the genetics of host response to commensal bacteria

**DOI:** 10.64898/2026.02.13.705705

**Authors:** Rémi Duflos, Fernando A. Rabanal, Rémy Zamar, Roxane Lion, Daniela Ramírez-Sánchez, Ana P. Leite Montalvão, Katrin Fritschi, Anette Habring, Natalie Betz, Joy Bergelson, Detlef Weigel, Fabienne Vailleau, Fabrice Roux

**Affiliations:** LIPME, INRAE, CNRS, Université de Toulouse, Castanet-Tolosan, France; Department of Molecular Biology, Max Planck Institute for Biology Tübingen, Tübingen, Germany; Center for Genomics and Systems Biology, Dept. of Biology, New York University, New York City, NY 10003, USA

**Keywords:** ecological genomics, microbiota, adaptive variation, fitness, *Arabidopsis thaliana*

## Abstract

**Background:** Microbiota members collectively contribute to individual performance in humans, animals, and plants. This led to the quest for probiotics to improve host health and reproductive performance. Although the efficacy of probiotics is known to be strongly affected by environmental factors, microbe-microbe interactions, and microbial strain identity, the effects of host genotype and the underlying genetic architecture have been overlooked. In addition, the evolutionary causes of such genetic variation are typically not addressed. In this study, we aimed to describe the genetic architecture of the adaptation of the host plant *Arabidopsis thaliana* to commensal bacterial members of its native microbiota by identifying candidate genes associated with fitness proxies and presenting signatures of natural selection.

**Results:** A Genome-Wide Association study conducted under field conditions revealed extensive variation within a new mapping population of 162 genotypes of *A. thaliana* scored for total seed production and its two underlying components, namely fruit number and mean seed number per fruit, in response to 13 commensal strains. In agreement with the strong ‘host genotype × commensal strain identity’ interactions observed for each reproductive trait, the polygenic genetic architecture was highly flexible among the 13 commensal strains. Candidate genes exhibited a significant enrichment in signatures of both local adaptation and balancing selection. In line with the phenotyped reproductive traits, we identified seven candidate genes with functions specifically and strongly linked to seed germination and fertility.

**Conclusions:** Our findings reveal the importance of genotype-by-genotype interactions when measuring fitness proxies on a wild plant species inoculated with key members of its native microbiota. In addition, this study improves our understanding of the genetic signatures of natural selection acting on native host-microbiota adaptive interactions.

## Background

Microbes play a central role in the uptake of nutrients, alleviation of abiotic stress, protection against pathogens, and modulation of reproductive functions, collectively contributing to individual performance in humans, wild and domesticated animals, wild plants, and crops [1–9]. This established relationship between members of the microbiota and individual performance has fueled a quest for interventions that rely on microbes. These can be probiotic live microorganisms that confer health benefits to the host or even microbes with therapeutic properties [10].

While probiotics and therapeutic microbes can improve physical and reproductive performances [11–14], prevent and manage diseases [10,15,16], or restore environmental health [14,17,18], their impact can be strongly affected by environmental factors including diet and climatic conditions [19–21], microbe-microbe interactions with members of the resident microbiota [22–24], and the microbial strain identity [24–26]. Quantitative Trait Loci (QTLs) that affect the structure of the host-associated microbiota have been identified [27,28], but studies of genetic variation in the host response to probiotic microbes are limited [26,29], focusing mostly on biomass and growth-related traits in plants [26], and usually not addressing the evolutionary causes of such genetic variation.

In this study, we aimed to describe the host genetic architecture of adaptation to individual inoculation with native commensal bacteria. To do so, we set up a Genome-Wide Association study (GWAS) in field conditions by inoculating a new regional mapping population of *Arabidopsis thaliana* from the southwest of France with 13 commensal bacterial strains isolated from the same geographical region. These 13 strains have been demonstrated to have a beneficial effect on the vegetative growth of a limited number of genotypes of *A. thaliana* in axenic conditions [30]. After fine-mapping down to the gene level QTLs associated with total seed production and its two underlying components, i.e., fruit number and mean number of seeds per fruit, we tested whether the candidate genes overlap significantly with complementary signatures of natural selection, namely local adaptation, selective sweeps, and balancing selection.

## Results

Fifty-four populations (each represented by three genotypes, hereafter called accessions) were chosen to represent both the genomic and ecological diversity among a set of 168 natural populations of *A. thaliana* located in the southwest of France [31–33] (Fig. S1, Dataset S1). The 162 accessions were inoculated in field conditions (Fig. S1) with 13 bacterial strains belonging to seven out of the 12 most abundant and prevalent commensal bacterial Operational Taxonomic Units (OTUs) of the phyllosphere of the original 168 *A. thaliana* populations [30,32]. Based on whole-genome sequencing, the seven OTUs were taxonomically affiliated with seven species: *Paraburkholderia fungorum* (OTU2), a new species from the *Oxalobacteraceae* family (OTU3a), a new species from the *Comamonadaceae* family (OTU4), *Pseudomonas moraviensis* (OTU5), *Pseudomonas siliginis* (OTU6), a new *Methylobacterium* species (OTU13), and a new bacterial species from the *Sphingomonadaceae* family (OTU29) [30]. Plants were inoculated with two strains per bacterial species, except for OTU4 for which only one strain was available [30]. Together with a non-inoculated control (i.e. mock treatment), this resulted in 14 treatments.

A total of 13,450 plants were phenotyped for three traits that have been demonstrated to be good proxies of reproductive performance in highly selfing species such as *A. thaliana* [34–37], i.e., number of fruits, mean fruit length that is highly correlated with the number of seeds per fruit [38], and the composite trait of total seed production, which was calculated from number of fruits × mean fruit length (Dataset S2). Extensive and significant genetic variation was observed among and within populations for total seed production and its two underlying components in the mock treatment and the 13 treatments with bacterial inoculation (Table S1, Dataset S3). Accordingly, the broad-sense heritability (*H*²) estimate was highly significant for each ‘trait × treatment’ combination (Dataset S4). The mean *H*² estimate across the 14 treatments was 0.63 for total seed production (min = 0.56, max = 0.69), 0.62 for fruit number (min = 0.54, max = 0.70), and 0.70 for mean fruit length (min = 0.62, max = 0.75), indicating that a very large fraction of phenotypic variance was explained by genetic variation among accessions under our field conditions (Dataset S4). Based on genotypic values, we then estimated the extent of the plant response of each *A. thaliana* accession to each bacterial strain relative to the mock treatment for each reproductive trait. We observed extensive variation among accessions for both the direction and the strength of plant response to inoculation for each reproductive trait (Fig. 1). For instance, across the 13 bacterial strains, the absolute mean plant response was 34.9% for total seed production (95^th^ percentile = 96.2%), 32.7% for fruit number (95^th^ percentile = 83.8%), and 8.6% for mean fruit length (95^th^ percentile = 23.2%) (Fig. 1*A,B,C*). While the mean plant response to inoculation was significantly dependent on the bacterial species, it was also strongly influenced by the strain identity within a given bacterial species, particularly for total seed production and fruit number (Fig. 1*A,B,C*, Table S2). Finally, that reaction norms often went into opposite directions for two strains of a given bacterial species is suggestive of strong and very specific genotype-by-genotype (G×G) interactions (Fig. 1*D,E,F*).

**Fig. 1.**
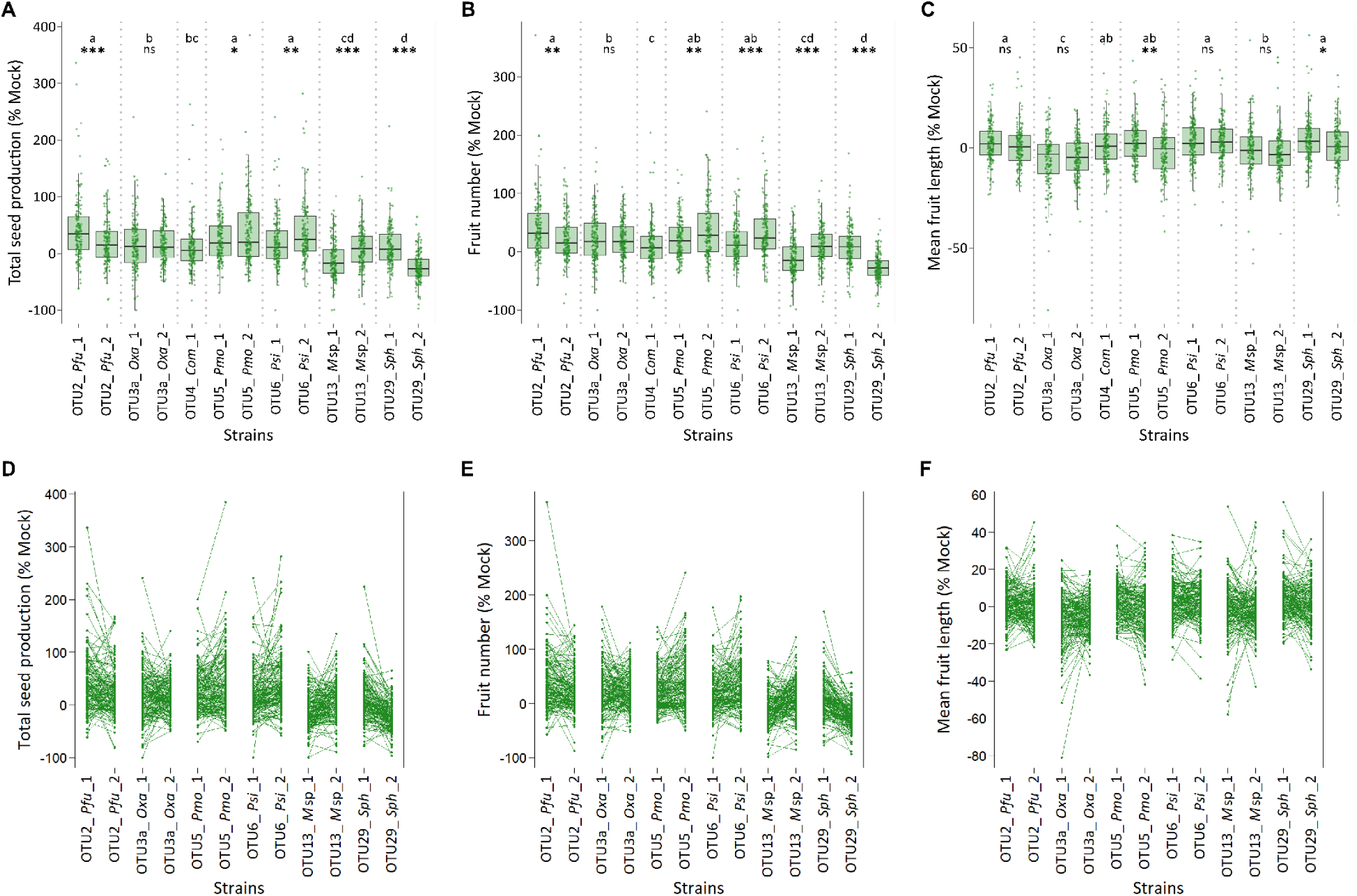
Phenotypic response of *A. thaliana* accessions to the individual inoculation of 13 commensal strains. Boxplots illustrating differences among bacterial species and between strains within a given bacterial species for total seed production (*A*), fruit number (*B*), and mean fruit length (*C*) expressed in percentage relative to the mock treatment. For each trait, different letters indicate different groups according to the treatments after a Ryan-Einot-Gabriel-Welsh (REGWQ) multiple-range test at *P* = 0.05. Differences between strains with a given bacterial species: *** *P* < 0.001, ** *P* < 0.01, * *P* < 0.05, ns: non-significant. Interaction plots illustrating the crossing reaction norms between strains within a given bacterial species for total seed production (*D*), fruit number (*E*), and mean fruit length (*F*) expressed in percentage relative to the mock treatment. Each *A. thaliana* accession is represented by a dot.

Importantly, for total seed production, we observed a strong and significant negative relationship between the plant response to inoculation with bacterial strains and the genotypic values in the mock treatment (Fig. 2*A*, Table S3). The inoculation with commensal strains had, on average, a positive effect on accessions with low total seed production in the mock treatment and a negative effect on accessions with high total seed production in the mock treatment (Fig. 2*A*). The strength of this negative relationship was significantly dependent on the bacterial species, but not on the strain identity within a given bacterial species (Table S3). For instance, the negative relationship was strong and highly significant for the response to OTU5 but weak and not significant for the response to OTU29 (Fig. 2*A*). A similar negative relationship was observed for fruit number (Fig. 2*B*, Table S3) and mean fruit length (Fig. 2*C*, Table S3).

**Fig. 2.**
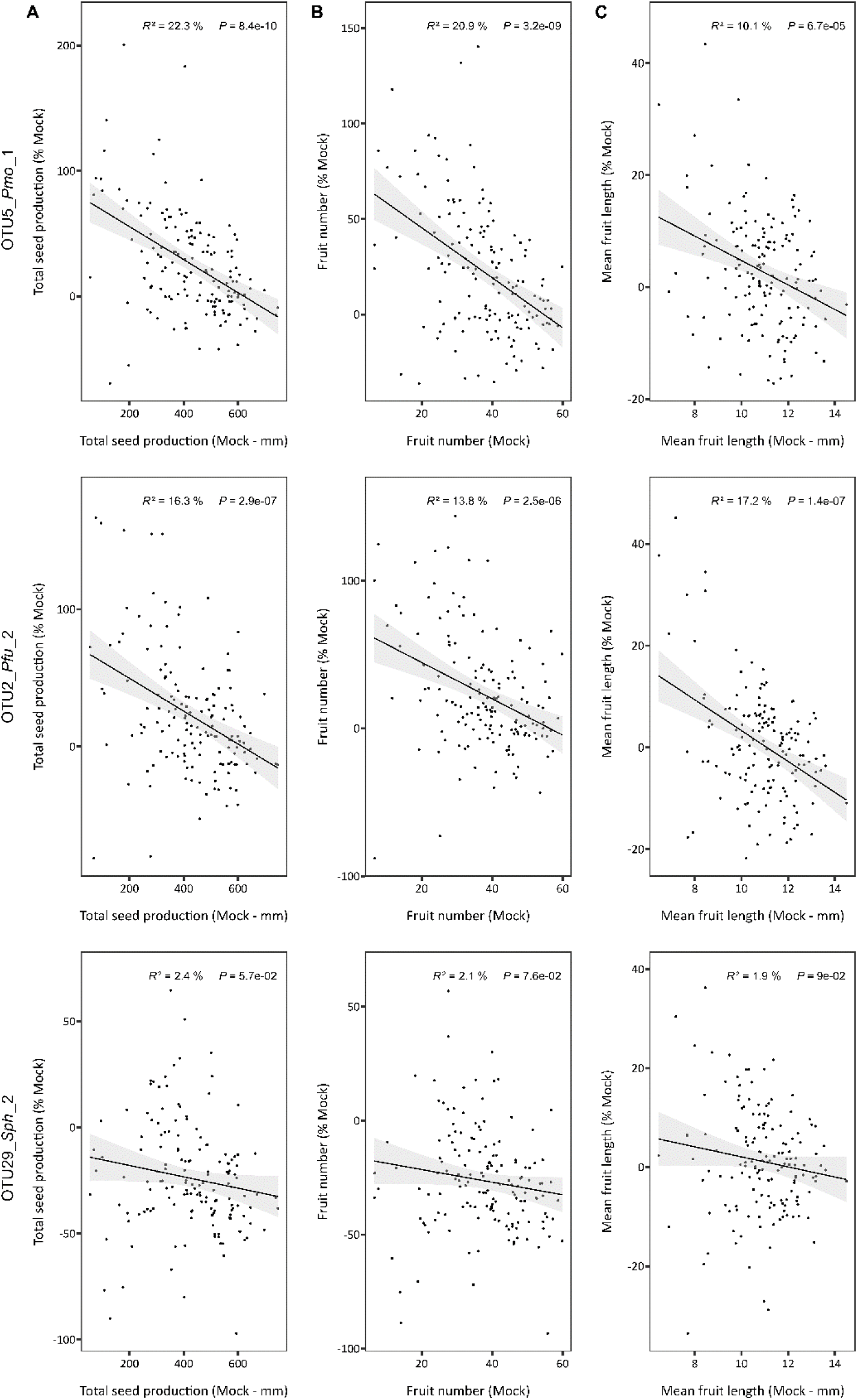
Negative relationships between the level of plant response to three commensal strains (expressed in percentage relative to the mock treatment) and the genotypic values in the mock treatment for total seed production (*A*), fruit number (*B*), and mean fruit length (*C*). *R*²: Percentage of the variance of plant response explained by the genotypic values in the mock treatment. Each *A. thaliana* accession is represented by a dot.

To enable the mapping of genotype-phenotype associations, we sequenced with the short-read sequencing technology Illumina 458 accessions from the southwest of France, including 146 accessions phenotyped in this study (Dataset S1). Population genomics analyses with 5,386,423 detected bi-allelic single nucleotide polymorphisms (SNPs) revealed that (i) genomic variation was weakly explained by geographic coordinates (*R*² = 8.9%), suggesting limited population structure, (ii) each accession was genetically different from each other, confirming that all 168 natural populations from the southwest of France are polymorphic [31], and (iii) linkage disequilibrium (LD) decays to *r*² = 0.5 within an average of around 0.75 kb (Fig. S2). To characterize the genetic architecture of plant response to inoculation with the 13 commensal bacterial strains, we combined a mixed model controlling for the effect of demographic history with a local score approach [39]. For each ‘reproductive trait × bacterial strain’ combination, the genetic architecture was highly polygenic with the detection of, on average, ∼21 QTLs per strain for total seed production, ∼23 QTLs per strain for fruit number, and ∼21 QTLs per strain for mean fruit length (Fig. 3*A*, Dataset S5). In agreement with the LD estimate, the average size of all 850 QTL intervals was only 3.3kb (median = 1.6kb, 5^th^ percentile = 0.2kb, 95^th^ percentile = 12.2kb) (Dataset S5). For each reproductive trait, the polygenic architecture was highly flexible among the bacterial strains, as illustrated by the L-shaped distribution of the degree of environmental pleiotropy (i.e. number of bacterial strains) of the candidate genes (Fig. 3*B*, Dataset S6).

**Fig. 3.**
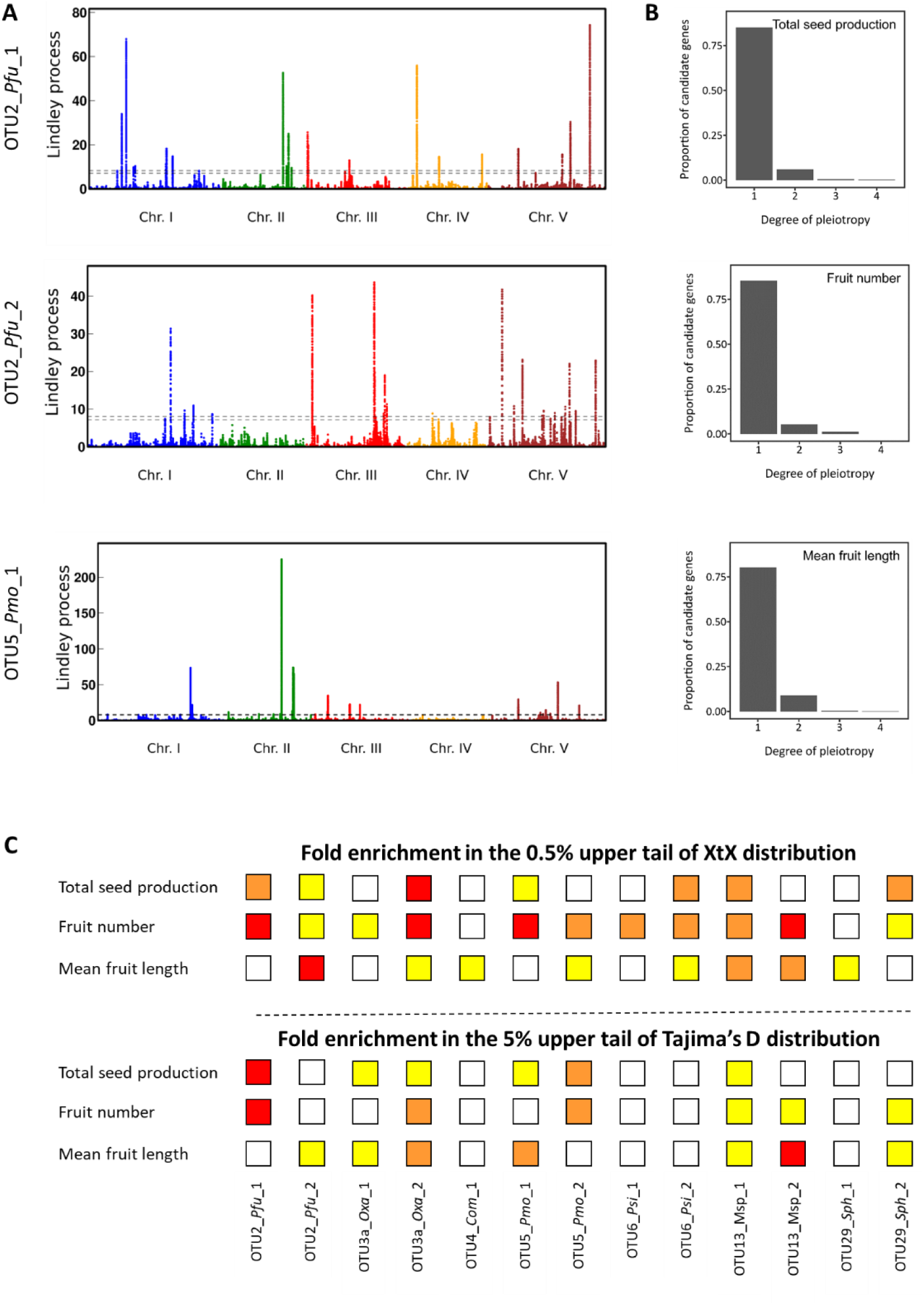
A genomic map of adaptation to commensal strains. (*A*) Manhattan plots of the Lindley process for the level of plant response to three commensal strains (expressed in percentage relative to the mock treatment) for total seed production. The dashed line indicates the chromosome-wide significance threshold. (*B*) Degree of environmental pleiotropy (i.e. number of strains) of candidate genes for each reproductive trait. (*C*) Significance of the fold enrichment in the 0.5% upper tail of XtX distribution (top panel) and the 5% upper tail of the Tajima’s D distribution (bottom panel) for each ‘phenotypic trait × bacterial strain’ combination. Red squares: *P* < 0.001, Orange squares: *P* < 0.01, Yellow squares: *P* < 0.05.

To determine whether the loci identified by GWA mapping had been shaped by natural selection, we first tested whether the SNPs in QTL intervals (hereafter called top SNPs) were enriched in a more general set of SNPs suggestive of adaptive spatial differentiation. To do so, we used a genome-wide selection scan previously obtained by estimating a Bayesian measure of spatial genetic differentiation (XtX) among *A. thaliana* populations from the southwest of France [31], with SNPs with the highest XtX values considered as the main candidates for signatures of local adaptation [40]. The 0.5% upper tail of the spatial genetic differentiation distribution was significantly enriched for top SNPs of two-thirds of the 39 ‘reproductive trait × bacterial strain’ combinations (Fig. 3*C*, Table S4). In a second approach, we tested whether the candidate genes significantly overlapped with signatures of selective sweeps. To do so, we identified genomic regions affected by recent positive selection, by performing a composite likelihood ratio (LR) test across the genome of the 458 Illumina whole-genome sequenced accessions [41]. Significant enrichment of GWA top SNPs in the upper tail of the LR distribution was rare for all 39 ‘reproductive trait × bacterial strain’ combinations (Dataset S7). In a third approach, we tested whether the candidate genes significantly overlapped with genes having signatures of balancing selection. To do so, we made use of telomere-to-telomere genome assemblies of 50 accessions from the southwest of France obtained by PacBio HiFi sequencing. Of these, 24 were previously released [42], and 26 were new (Dataset S8). We then estimated Tajima’s D values for 36,330 nuclear gene models lifted over from the TAIR10 reference genome (Dataset S9). Tajima’s D values followed a right-skewed distribution (mean value = -0.283, median value = -0.472) (Fig. S3, Dataset S9). The 5% upper tail of Tajima’s D distribution (suggestive of balancing selection) was significantly enriched for at least one reproductive trait for 10 of the 13 bacterial strains (Fig. *3C*, Table S5).

In agreement with the L-shaped distribution of the degree of environmental pleiotropy, 46 of the 55 candidate genes with both signatures of local adaptation and signatures of balancing selection were specific to one of our 13 bacterial strains (Dataset S6). One such example was the gene *STRESS INDUCED FACTOR 1* (*AT1G51840*), encoding for a kinase-like protein (Fig. 4*A,D*). For the nine candidate genes relevant for infection with more than one bacterial strain, similar effects were often observed for the plant response to two bacterial strains between highly divergent haplotypes (Dataset S6). Examples here include the gene *AGAMOUS-LIKE 83* (*AT5G49490*) encoding for a MADS box transcription factor for total seed production (Fig. S4) and the gene U2B’’ (*AT2G30260*) encoding for a small nuclear ribonucleoprotein for mean fruit length (Fig. 4*B,E*). U2B’’ is a direct regulatory target of MYC2 [43], a protein involved in a microbiota–root–shoot circuit that has been shown to enhance plant growth [44]. We also identified a few candidate genes for which the phenotypic effect between two highly divergent haplotypes can be reversed between ‘reproductive trait × bacterial strain’ combinations (Dataset S6). For instance, the differential effect of OTU5_*Pmo*_1 inoculation on fruit number between the two haplotypes of the gene EMBRYO DEFECTIVE 1974 (*AT3G07060*), which encodes for an NHL domain-containing protein, is reversed for mean fruit length in response to OTU2_*Pfu*_2 (Fig. 4*C,F*).

**Fig. 4.**
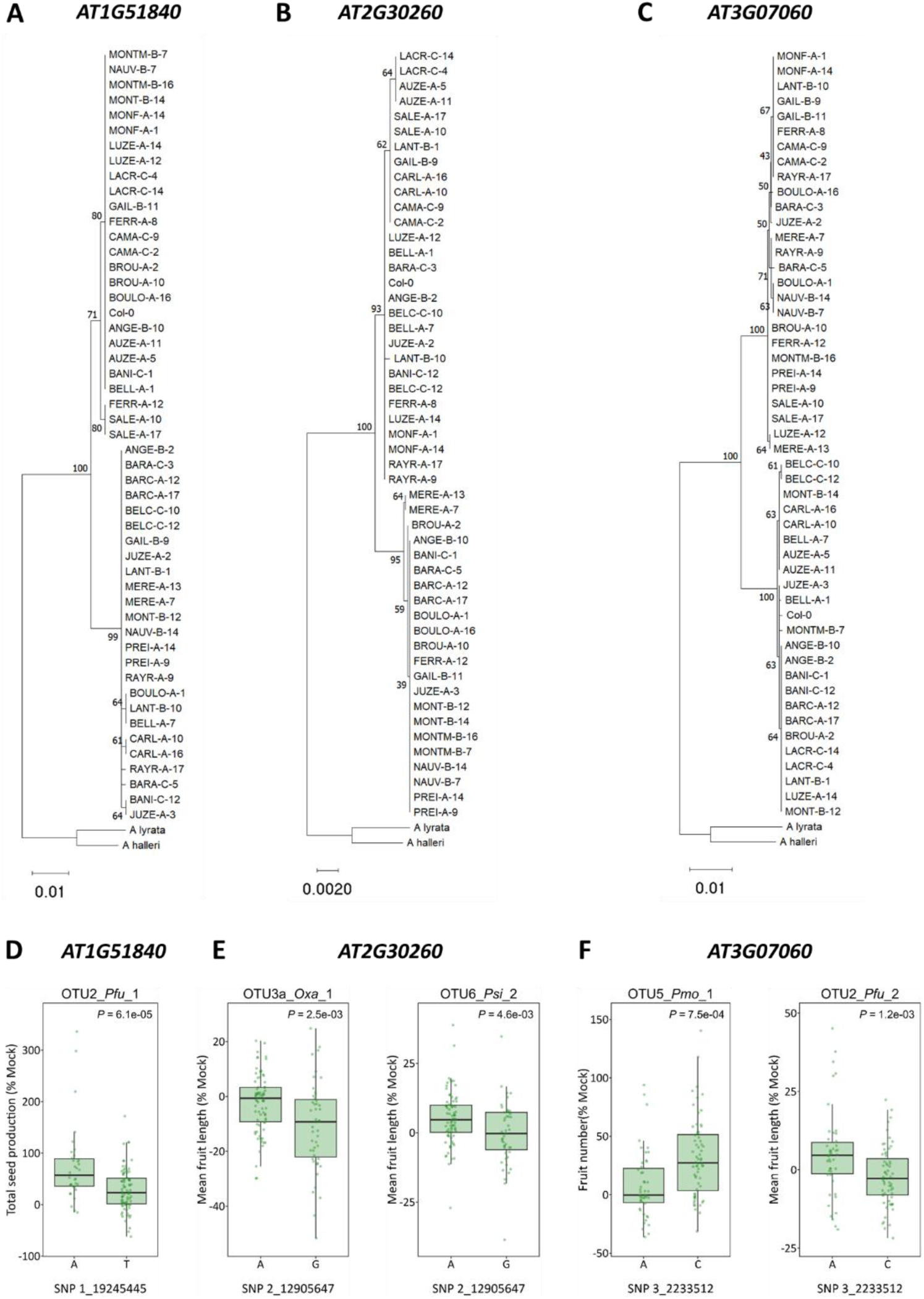
Examples of candidate genes presenting both signatures of local adaptation and signatures of balancing selection. (*A*), (*B*) and (*C*) Neighbor-joining tree of three candidate genes. (*D*) Plant response for total seed production to the commensal strain OUT2_*Pfu*_1 according to the two main haplotypes of *AT1G51480* represented by the top SNP 1_19245445. (*E*) Plant response for mean fruit length to two commensal strains according to the two main haplotypes of *AT2G30260* represented by the top SNP 2_12905647. (*F*) Contrasted plant response to two commensal strains according to the two main haplotypes of *AT3G07060* represented by the top SNP 3_2233512. (*D*), (*E*) and (*F*) Each *A. thaliana* accession is represented by a dot.

We next looked for biological processes significantly overrepresented among the candidate genes identified for each reproductive trait by comparing them to the overall class frequency in the *A. thaliana* MapMan annotation. We identified two classes that were significantly enriched in the three lists of unique candidate genes, ‘RNA’ and ‘signaling’ (Fig. S5, Dataset S10), consistent with the main biological pathways identified using lab-induced mutations and through previous GWAS results for microbiota regulation in *A. thaliana* [28].

To investigate the molecular functions involved in the adaptive response of *A. thaliana* to inoculation with these commensal bacterial strains, we focused on candidate genes identified for total seed production, belonging to enriched biological processes, and with signatures of both local adaptation and balancing selection. Based on these criteria, we identified seven candidate genes that are either involved in abiotic stress responses (e.g., oxidative stress, drought) or closely associated with hormonal regulation processes (e.g., auxin, abscisic acid), seed germination, and embryo development (Table S6). Consistent with the measure of a reproductive trait, the seven candidate genes exhibit specific or elevated expression levels in reproductive organs, including seeds and floral organs, but not pollen (Table S6).

## Discussion

While almost 300 GWA studies have been conducted to describe the genetic architecture of plant resistance to pathogens [45], there are surprisingly few reports on the genetic architecture of plant response to commensals [26], even though commensals represent a much larger fraction of the microbiota than pathogens. Here, we characterized the genetic architecture underlying the adaptive response of *A. thaliana* to the inoculation of commensal members of its native bacterial microbiota. We also identified candidate genes associated with fitness-related traits and presenting signatures of natural selection, including signatures of local adaptation and balancing selection.

Under our field conditions, there was an extensive variation among *A. thaliana* accessions for the level of response, both positive and negative, to a given commensal strain. The strongest positive responses were comparable to the percentage of yield increments typically seen in certain ‘crop variety × Plant Growth-Promoting Bacteria (PGPB)’ combinations [14]. Together with the detection of strong genotype-by-genotype interactions, this suggests that whether a commensal strain is considered a PGPB might be highly dependent on the host genotype tested. If similar observations are made in humans and animals in response to probiotics, this calls for the development of personalized approaches to probiotic treatment in both medicine and agriculture.

Previous studies have reported common genetic and molecular mechanisms in plant host response to pathogens and commensals [28,46–48]. Our study confirms the importance of ‘signaling’ genes involved in plant innate immunity, including kinase-related genes, in mediating host response to inoculation with bacterial commensals. We also identified a category of candidate genes that have seldom been reported as mediating plant-microbe interactions, namely genes with elevated expression levels in reproductive organs. This new category of candidate genes may originate from focusing on reproductive rather than vegetative traits.

As previously observed on disease resistance genes [49], we have shown how three complementary methods for estimating adaptive values (i.e., fitness assays, population genomics, and molecular evolution) yield information on the evolutionary and ecological forces acting on polymorphic genes involved in host-microbiota native interactions. The enrichment in signatures of balancing selection is in agreement with the negative relationship observed at the phenotypic level. Interestingly, the observed balancing selection might be reinforced by contrasting phenotypic effects on fitness-related traits in response to multiple commensal strains, leading to highly divergent haplotypes. Because hosts are rarely exposed to a single commensal species, a better understanding of coevolutionary quantitative genetics of native host-microbiota adaptive interactions would benefit from exploring diffuse biotic interactions rather than tightly associated host–commensal pairs.

## Materials and Methods

### Plant and bacterial material

For the 162 accessions used in this study, seeds from maternal plants sampled in natural populations were collected in May 2015. Differences in the maternal effects among the accessions were reduced by growing one plant of each accession for one generation [50]. The 13 bacterial strains considered in this study were isolated in 2020 from *A. thaliana* rosettes sampled in 2015 in local populations in the southwest of France [30] and were stored in a 20% glycerol solution at -80°C.

### Experimental design, growth conditions, and inoculation procedure

The experimental design, growth conditions, and inoculation procedure have been extensively detailed in [50]. Briefly, a field experiment of 15,552 plants was set up at the INRAE center of Toulouse (France) using a split-plot design arranged as a randomized complete block design (RCBD) with 16 treatments nested within six experimental blocks (supplementary fig. S1). The 16 treatments correspond to two mock treatments and the individual inoculation of 14 bacterial strains, including the 13 commensal strains considered in this study and the strain JACO-CL of the bacterial pathogen *Pseudomonas viridiflava* (OTU8) [51] that was considered for another study. Seeds were sown on 18 March 2021.

Bacterial strains were grown on solid medium in Petri dishes (TSA for OTU5, OTU6, and OTU8; TSB for OTU2; R2A for OTU3, OTU4, OTU13, and OTU29). On 14 April 2021, when most plants reached a 5-6 leaf stage, a volume of 1 mL of bacterial inoculum (OD_600 nm_ of 0.01) was dispensed on each rosette. A volume of 1mL of sterile water was dispensed on each rosette of the plants of the two mock treatments. To facilitate the penetration of bacteria cells into plant organs, the Tween® 20 surfactant was added to each inoculum at a final concentration of 0.01%.

### Phenotyping

Three reproductive traits were measured on plants that were collected after completing their life cycle on 17-18 June 2021. Following [36,38,52], fruit number corresponds to the total number of fruits produced on the primary shoot, the primary branches on the primary shoot, and the basal branches. Mean fruit length corresponds to the average length of three representative fruits. Total seed production corresponds to fruit number multiplied by mean fruit length.

### Statistical analysis

To estimate the natural genetic variation of the three reproductive traits among and within populations for each treatment (mock + inoculation with 13 bacterial strains), the following linear model (function *lm* under the R environment) was used:

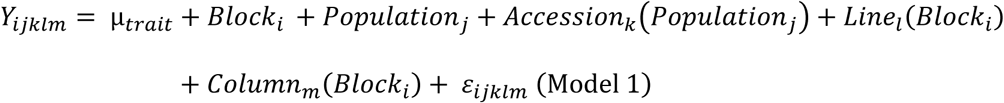

where Y is one of the three phenotypic traits, µ is the overall mean of the phenotypic data, ‘Block’ accounts for differences in micro-environmental conditions among blocks, ‘Line(Block)’ and ‘Column(Block)’ accounts for difference in micro-environmental conditions within blocks, ‘Population’ corresponds to the genetic differences among the 54 populations, ‘Accession’ accounts for mean genetic differences between the three accessions within populations, and ‘ε’ is the residual term. All the factors were treated as fixed effects because the levels of no factor were the random samples from a population to which we intended to extrapolate.

For each of the 42 ‘phenotypic trait × treatment’ combinations (*i.e*., three traits × 14 treatments), genotypic values of the accessions were estimated by calculating least-squares (LS) mean values of the ‘Accession’ term in the following linear model (function *lm* under the R environment):

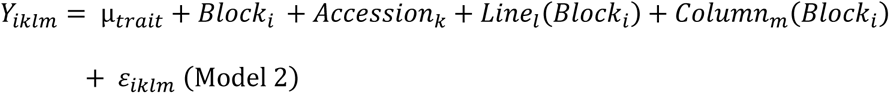

We then applied the function *lsmeans* (library *lsmeans* under the R environment) on Model 2. With the ‘Block’ and ‘Accession’ factors treated as random effects, the percentage of phenotypic variance explained by the ‘Block’ and ‘Accession’ terms was then estimated by applying the function *varcorr* (library lme4 under the R environment) on model 2. Following [53], *H*² values were estimated using the following formula:

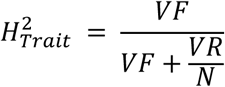

where ‘*VF*’ corresponds to the genetic variance among the accessions, “*VR*” is the residual variance, and ‘*N*’ is the mean number of biological replicates per accession (N = 6 in this study).

Based on genotypic values, we estimated for each reproductive trait the extent of plant response of each accession to each strain using the following formula:

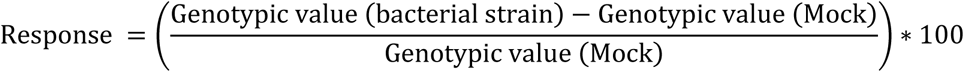

For each trait, the estimated plant responses were then used to (i) compare phenotypic variation among the 13 treatments with a bacterial strain, (ii) study the relationship between plant response to a specific bacterial strain and genotypic values observed in the mock treatment, and (iii) run GWA mapping analyses (see below).

### Short-read sequencing: DNA extraction, library preparation, sequencing, bioinformatic analyses, patterns of linkage disequilibrium, and population structure

We sowed 503 *A. thaliana* accessions from the southwest of France on Jiffy pots, including the 162 accessions used in this study. Of these, 475 accessions were germinated and grown under controlled conditions (20°C, 16 h photoperiod). Total DNA was extracted from rosette samples using a modified version of the protocol developed to extract high molecular weight gDNA [54]. DNA was quantified by PicoGreen, and libraries were constructed using a Nextera protocol modified to include smaller volumes [55,56]. Library molecules were size selected on a Blue Pippin instrument (Sage Science, Beverly, MA, USA), and libraries were sequenced on an Illumina Nextseq2000 using a paired-end read length of 2×150 pb (Genome Center, Max Planck Institute for Biology, Tübingen, Germany).

Raw reads of each of the 475 accessions were mapped onto the TAIR10 *A. thaliana* reference genome Col-0 with BWA (v.0.7.17)[57]. After removing 17 accessions with a very low coverage depth, a semi-stringent SNPCalling across the genome was performed for each of the 458 remaining accessions (including 146 accessions phenotyped in this study) with SAMtools mpileup (v.1.19)[58] and VarScan (v.2.5.2)[59] with the parameters corresponding to a theoretical sequencing coverage of 30X and the search for homozygous sites. A set of 5,386,423 bi-allelic SNPs was detected across the 458 accessions. Considering only SNPs with a minor allele relative frequency (MARF) > 0.1, the extent of linkage disequilibrium within 30-kb windows on each chromosome was estimated using VCFtools (v.0.1.16) [60]. A linkage pruning was performed with PLINK (v.1.9)[61] using the following parameters “--set-missing-var-ids @:# --indep-pairwise 50 10 0.1”. A PCA was then run on the resulting bi-allelic SNP matrix (N = 1,210,830 SNPs) using the following parameters “--set-missing-var-ids @:# --extract for_pca.prune.in --make-bed –pca”.

### Long-read sequencing: DNA extraction, library preparation, sequencing, genome assembly and annotation

Fifty *A. thaliana* accessions from the southwest of France were grown at 23°C under 16 hours of light, and 300 mg of tissue powder from 21-day-old rosettes from multiple plants, or from single 26-day-old individual plants, was used for high-molecular-weight (HMW) genomic DNA extraction. The methods for HMW-DNA extraction, PacBio HiFi library preparation, genome assembly using hifiasm (version 0.16.1-r375) [62], and quality assessment were extensively detailed in the previous release of 24 of these genome assemblies [42]. An additional 26 genome assemblies, processed using the same methods, are newly released as part of this study (Dataset S8). The annotation of protein-coding genes was carried out with Liftoff (version 1.6.2) [63] (‘-copies -sc 0.98 -polish - cds’), using as input annotation file ‘Araport11_GFF3_genes_transposons.201606.gff’ downloaded from https://www.arabidopsis.org.

### GWA mapping combined with a local score approach (GW-LS)

GWA mapping was run using a mixed model approach implemented in the software EMMAX (Efficient Mixed-Model Association eXpedited) [64]. This model includes a genetic kinship matrix as a covariate to control for the effect of the demographic history of the 54 populations from the southwest of France. In this study, we discarded SNPs with more than 96 missing values across the 146 whole-genome sequenced accessions phenotyped in this study, and we only considered the SNPs with a minor allele relative frequency (MARF) >= 10%, leaving us with 630,269 SNPs.

A local score approach [39] with a tuning parameter ξ = 2 was applied to the *p* values generated by EMMAX. Significant SNP-phenotype associations were identified by estimating a chromosome-wide significance threshold for each chromosome.

### Enrichment analyses

To test whether the top SNPs exhibit signatures of local adaptation, we followed the method previously described in [52] by calculating a fold enrichment of the top SNPs in the extreme upper tail of the XtX distribution obtained for the set of 168 natural populations of *A. thaliana* [31]. For a given SNP, XtX is a measure of the variance of the standardized population allele frequencies, which results from a rescaling based on the covariance matrix of population allele frequencies [40]. Following the methodology described in [65], the statistical significance of enrichment was assessed by running 10,000 null circular permutations across the five chromosomes of *A. thaliana*.

Genomic regions affected by recent positive selection were identified by performing a composite likelihood ratio (LR) test implemented in SweepFinder2 [41] across the genome of the 458 Illumina whole-genome sequenced accessions. LR tests were computed for four window sizes (1kb, 5kb, 10kb, and 50kb). To test whether the candidate genes exhibit signatures of selective sweeps, we calculated a fold enrichment of the candidate genes in the extreme upper tail of the LR distribution. Using a custom Python script (v.3.9.2), the statistical significance of enrichment was assessed by running across the five chromosomes of *A. thaliana* 10,000 null permutations, considering the block structure of the candidate genes underlying the QTLs.

Tajima’s D values were calculated using a homemade Python script (v.3.9.0). We only considered the 37,009 nuclear genes annotated in Araport11 with a full-length genomic sequence (including the 5’ and 3’ regions) > 100bp. The full-length genomic sequence of each gene was blasted on the genome assemblies of the 50 accessions from the southwest of France using NCBI BLAST+ tool (v.2.12.0) [66]. For each gene, we retrieved hits with a sequence-identity cutoff of 60% over 50% of the sequence. An absence of hits for at least one accession was found for 4,011 genes, suggesting presence/absence polymorphisms. When multiple hits were retrieved, the best hit was kept. For each gene, sequences were aligned using MAFFT (v.7.520) [67]. All indels were then removed from the final alignment and Tajima’s D was calculated following the formula in [68]. To test whether the candidate genes exhibit signatures of balancing selection, we calculated a fold enrichment of the candidate genes in the extreme upper tail of Tajima’s D distribution. The statistical significance of enrichment for signatures of balancing selection was assessed as previously described above for signatures of selective sweeps.

Using a custom Python script, we retrieved the candidate genes underlying the QTL intervals for each of the three reproductive traits. To identify biological pathways significantly over-represented (*P* < 0.05), the lists of the candidate genes were then submitted to the classification SuperViewer tool on the University of Toronto website (https://bar.utoronto.ca/ntools/cgi-bin/ntools_classification_superviewer.cgi) using the MapMan classification.

### Phylogenetic trees

Phylogenetic trees were built using MEGA (v.11.0.13)[69]. Each phylogenetic tree was rooted with orthologous sequences of the candidate gene retrieved from the genomes of *Arabidopsis lyrata* [70] and *Arabidopsis halleri* (https://phytozome-next.jgi.doe.gov/info/Ahalleri_v2_1_0). The number of bootstrap replications was 1,000. The trees were drawn to scale, with branch lengths in the same units as those of the evolutionary distances used to infer the phylogenetic tree. The evolutionary distances were computed using the Maximum Composite Likelihood method and are in the units of the number of base substitutions per site.

## Data availability

The raw Illumina sequencing data used for this study are available at the NCBI Sequence Read Archive (http://ncbi.nlm.nih.gov/sra) through the study accession SRP532425. The genome assemblies and the corresponding HiFi long-reads used in this study are available at the European Nucleotide Archive (https://www.ebi.ac.uk/ena/browser/home) under accession numbers PRJEB80227 (26 assemblies released in this study) and PRJEB55353 (24 assemblies previously released) [42]. The phenotypic data sets are available as supplementary Datasets.

## Acknowledgments

We thank Choghag Demirjian, Chrystel Gibelin-Viala, and Paul Baumet for their assistance during plant harvesting. We thank Kevin Murray for helpful discussions on the Illumina sequencing of the genomes of the 458 accessions of *A. thaliana*.

## Author Contributions

R.D., F.V., and F.R. designed research. R.D., F.A.R., R.Z., R.L., D.R.S., A.P.L.M., K.F., A.H., and N.B. performed research. R.D. and F.A.R. analyzed data. R.D., F.A.R., J.B., D.W., F.V., and F.R. wrote the paper.

## Declarations

### Ethics approval and consent to participate

not applicable.

### Consent for publication

not applicable.

## Supplementary Datasets

**Dataset S1.** Genetic and genomic resources associated with 503 accessions collected from 168 natural populations of *Arabidopsis thaliana* located in the southwest of France. **Dataset S2.** Raw data for the three reproductive traits scored on 14,580 plants in a field experiment conducted at INRAE Toulouse (France).

**Dataset S3.** Genotypic values of three reproductive traits for the accessions grown in field conditions in 14 treatments (mock treatment + 13 treatments with commensal strains).

**Dataset S4.** Basic statistics related to genotypic values and broad-sense heritability values (*H*²) for the three reproductive traits scored in field conditions in 14 treatments (mock treatment + 13 treatments with commensal strains).

**Dataset S5.** Genetic architecture of the three reproductive traits in the 13 treatments with commensal strains.

**Dataset S6.** Degree of environmental pleiotropy for each of the candidate genes identified by combining a mixed model with a local score approach.

**Dataset S7.** Enrichment in signatures of selective sweeps.

**Dataset S8.** Summary of the assembly characteristics of the 50 whole-genome sequenced accessions from the southwest of France.

**Dataset S9.** Tajima’s D values for 36,330 nuclear genes.

**Dataset S10.** For each reproductive trait, a list of candidate genes belonging to enriched biological processes.

