## Supplementary Information for "Natural selection acting on the genetics of host response to commensal bacteria"

**This PDF file includes:**

Supplementary Figures 1 to 5

Supplementary Tables 1 to 6

**Supplementary Fig. 1.** Mapping population from the southwest of France and field experiment. (*A*) Location of the 168 natural populations of *A. thaliana* in the southwest of France. Blue dots correspond to the 54 populations phenotyped in field conditions. The red dot corresponds to the field station at the INRAE center of Toulouse (France). (*B*) Illustration of the field experiment conducted at the INRAE center of Toulouse (France).

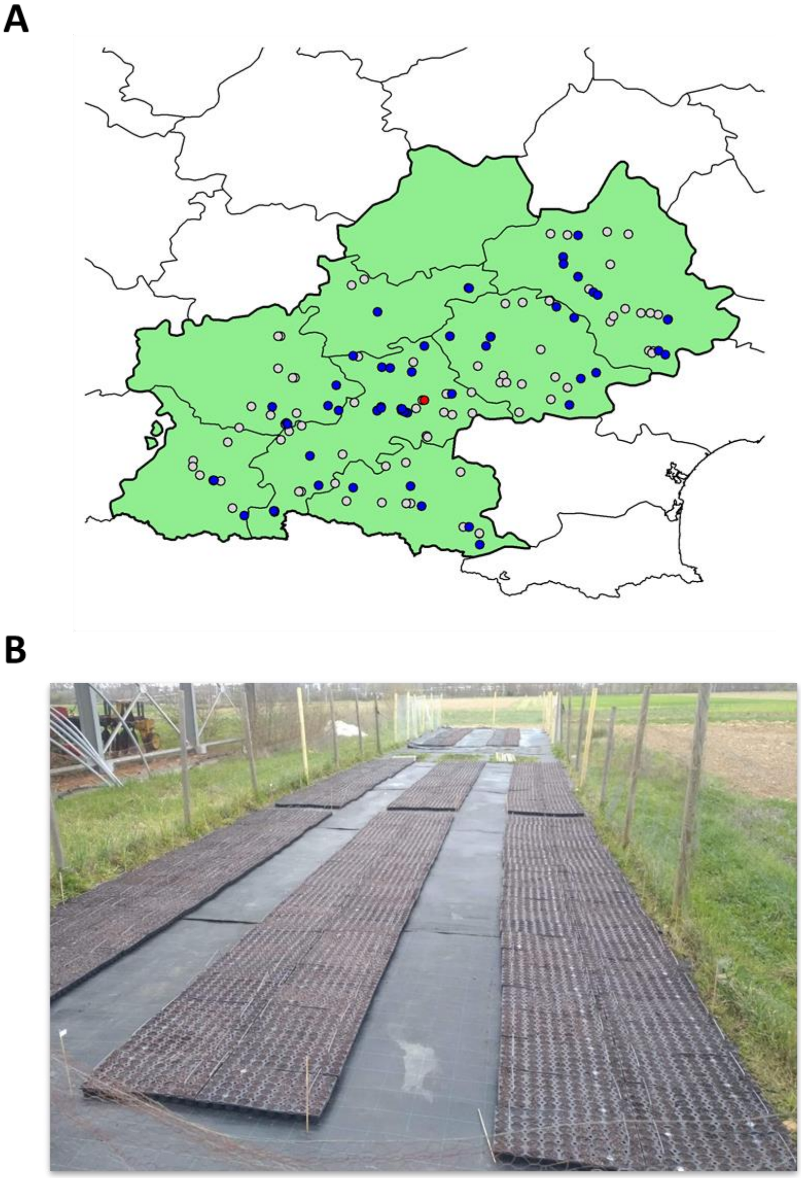

**Supplementary Fig. 2.** Genomic patterns of the mapping population of 458 accessions from the southwest of France. (*A*) Genomic space represented by the first two axes of a Principal Component Analysis run on the SNP matrix. Dots correspond to the 458 natural accessions. (*B*) Distribution of kinship coefficients. Only 0.84% of accession pairs have a kinship coefficient > 0.95. The pairwise genetic distance matrix is weakly explained by latitude, longitude, and elevation (PERMANOVA, latitude: $F = 10.3$, $P < 0.001$; longitude: $F = 9.2$, $P < 0.001$; elevation: $F = 6.9$, $P < 0.001$; latitude × longitude: $F = 5.4$, $P < 0.001$; latitude × elevation: $F = 5.4$, $P < 0.001$; longitude × elevation: $F = 4.8$, $P < 0.001$; latitude × longitude × elevation: $F = 4.1$, $P < 0.001$; $R^2 = 8.9\%$). (*C*) Decay of linkage disequilibrium ($r^2$) with physical distance.

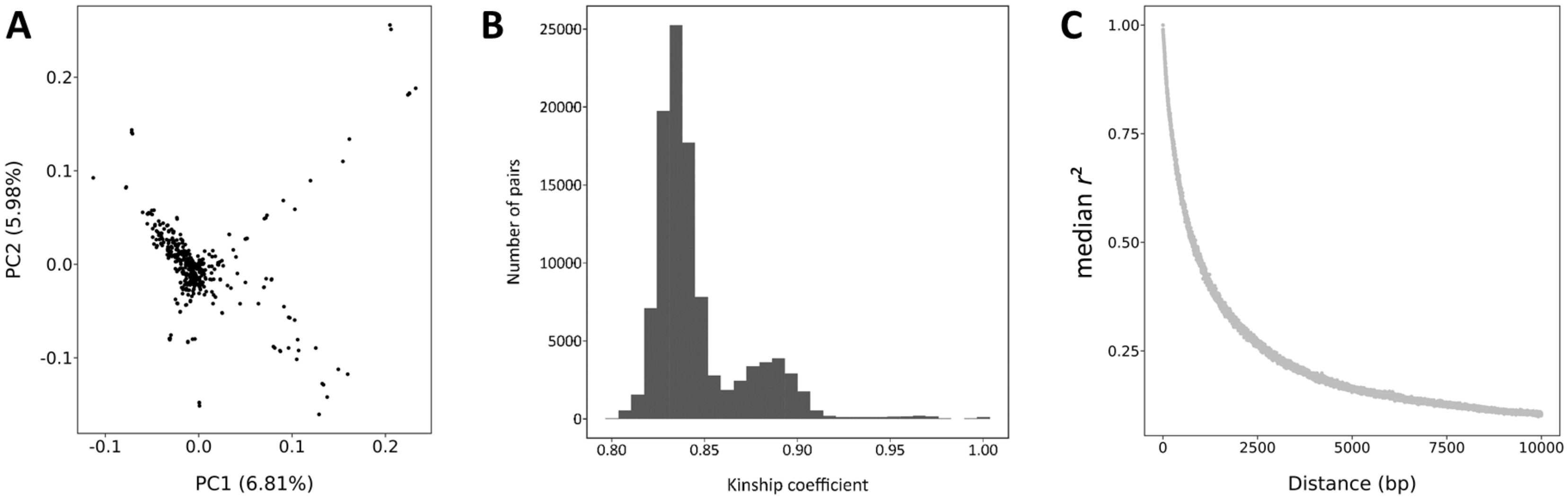

**Supplementary Fig. 3.** Distribution of Tajima's D values across 36,330 genes in 50 accessions from the southwest of France.

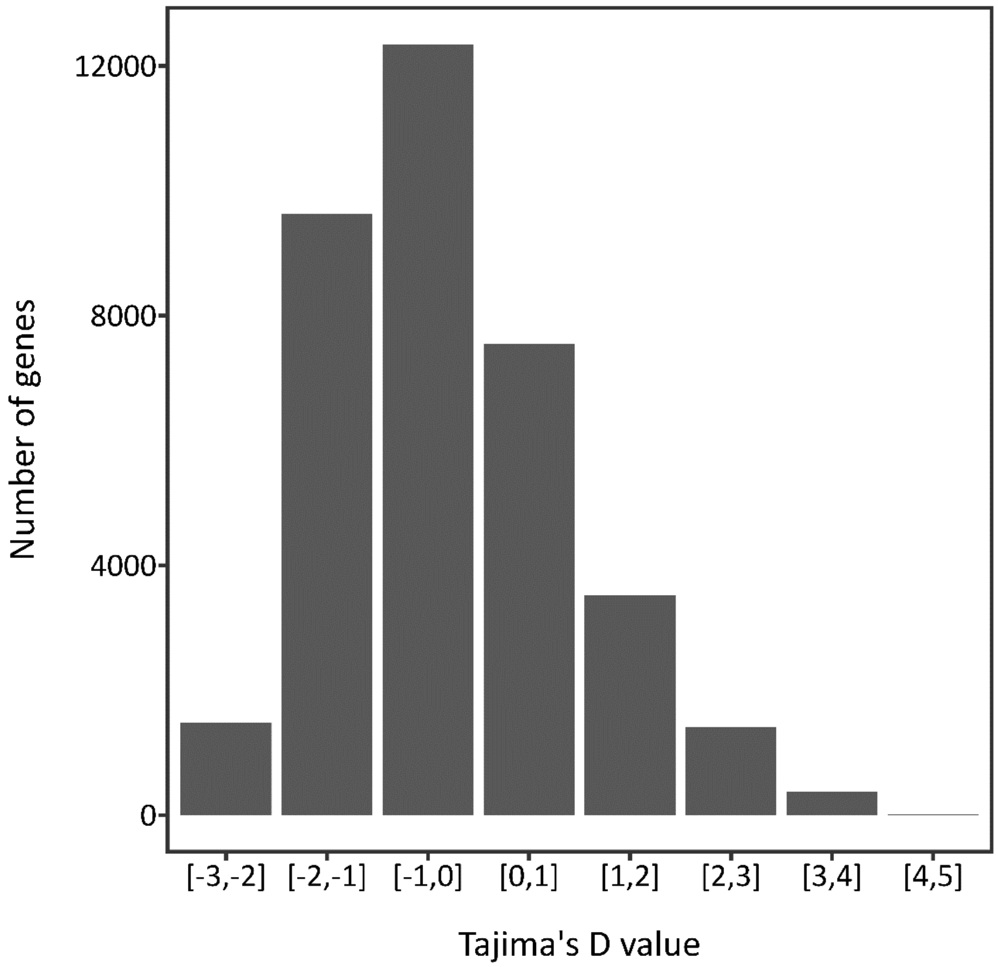

**Supplementary Fig. 4.** The candidate gene *AT5G49490* presenting signatures of local adaptation and signatures of balancing selection. (*A*) Neighbor-joining tree. (*B*) Plant response for total seed production to two commensal strains according to the main haplotypes of *AT5G49490*.

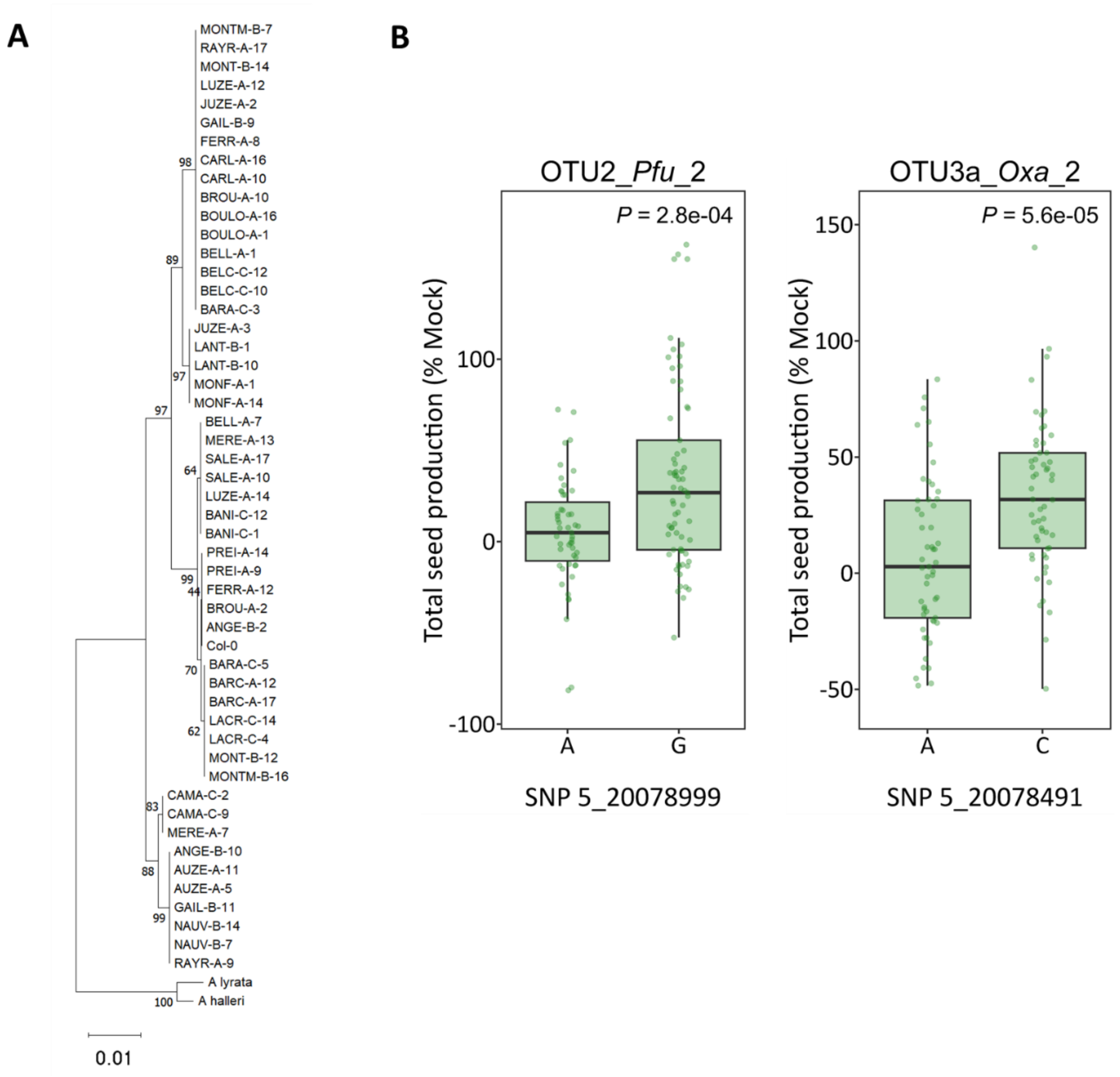

**Supplementary Fig. 5.** Enriched biological processes in response to the 13 bacterial strains.

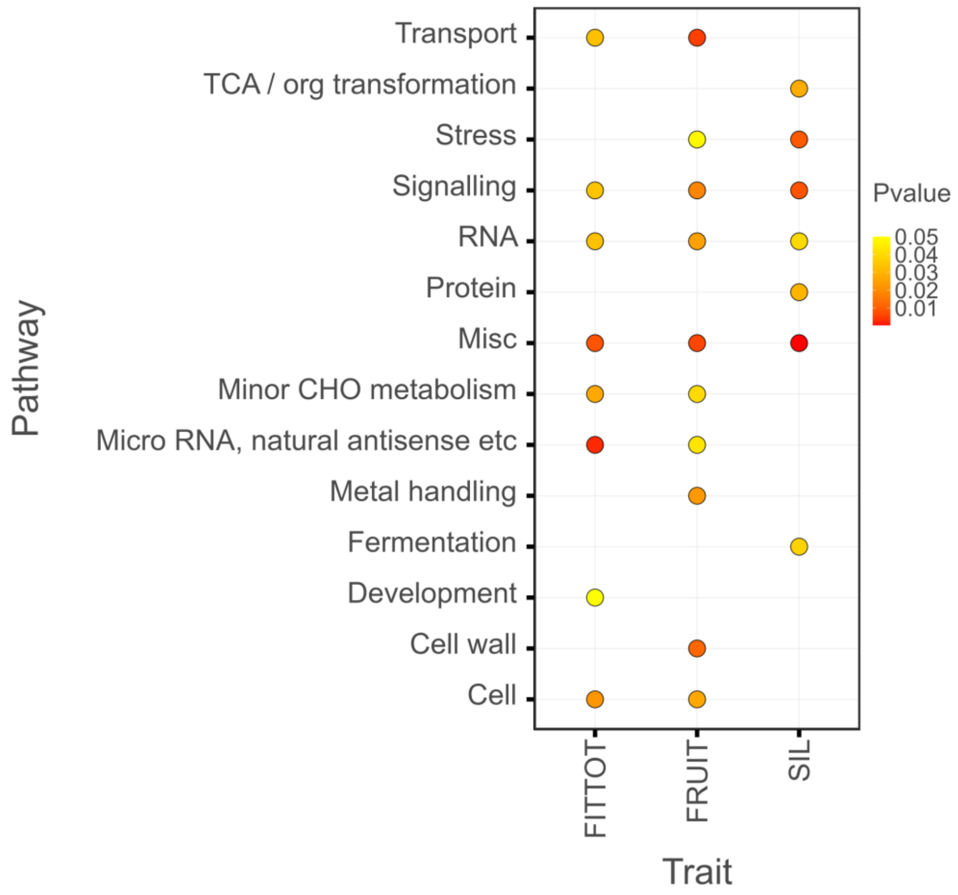

**Supplementary Table 1.** Genetic variation among and within populations for each reproductive trait in the 14 treatments (mock treatment + 13 treatments with bacterial strains).

| | | Total seed production | | Fruit number | | Mean fruit length | |
|---|---|---|---|---|---|---|---|
| Treatment | Model terms | *F* | *P* | *F* | *P* | *F* | *P* |
| Mock | Block | 57.4 | 9.36E-55 | 60.9 | 7.03E-58 | 5.5 | 4.92E-05 |
| | Population | 8.9 | 3.04E-58 | 8.7 | 4.12E-56 | 11.3 | 7.64E-76 |
| | Accession(Population) | 1.8 | 3.00E-06 | 1.8 | 5.12E-06 | 2.4 | 1.38E-11 |
| | ((Line(Mock_replicate))Block) | 11.7 | 6.02E-23 | 12.2 | 6.92E-24 | 3.6 | 2.48E-05 |
| | ((Column(Mock_replicate))Block) | 11.6 | 9.74E-23 | 11.8 | 3.61E-23 | 4.5 | 3.16E-07 |
| OTU2_*Pfu* _1 | Block | 14.6 | 1.23E-13 | 16.7 | 1.43E-15 | 4.2 | 9.99E-04 |
| | Population | 4.3 | 2.12E-19 | 4.0 | 1.23E-17 | 5.6 | 6.63E-28 |
| | Accession(Population) | 1.8 | 9.77E-06 | 1.8 | 1.02E-05 | 1.8 | 2.98E-05 |
| | Line(Block) | 4.9 | 6.27E-05 | 5.0 | 4.65E-05 | 1.1 | 3.47E-01 |
| | Column(Block) | 2.8 | 1.15E-02 | 2.2 | 4.47E-02 | 1.1 | 3.54E-01 |
| OTU2_*Pfu*_ 2 | Block | 16.3 | 3.28E-15 | 19.7 | 2.82E-18 | 6.9 | 2.88E-06 |
| | Population | 4.2 | 8.20E-19 | 3.8 | 3.21E-16 | 4.7 | 5.52E-22 |
| | Accession(Population) | 1.7 | 1.87E-04 | 1.7 | 4.68E-05 | 1.6 | 1.05E-03 |
| | Line(Block) | 5.9 | 5.35E-06 | 8.7 | 4.27E-09 | 1.9 | 8.06E-02 |
| | Column(Block) | 0.5 | 7.87E-01 | 0.6 | 7.01E-01 | 0.6 | 7.67E-01 |
| OTU3_*Oxa* _1 | Block | 42.6 | 3.98E-38 | 47.0 | 1.11E-41 | 1.7 | 1.42E-01 |
| | Population | 4.6 | 9.75E-22 | 4.6 | 1.26E-21 | 5.2 | 7.89E-25 |
| | Accession(Population) | 1.8 | 7.53E-06 | 1.8 | 8.99E-06 | 1.7 | 1.49E-04 |
| | Line(Block) | 3.7 | 1.35E-03 | 5.9 | 4.60E-06 | 1.2 | 3.17E-01 |
| | Column(Block) | 1.1 | 3.87E-01 | 1.2 | 3.19E-01 | 2.3 | 3.65E-02 |
| OTU3_*Oxa* _2 | Block | 12.0 | 3.73E-11 | 13.8 | 7.99E-13 | 6.0 | 1.84E-05 |
| | Population | 4.2 | 1.11E-18 | 4.3 | 1.73E-19 | 4.1 | 2.77E-18 |
| | Accession(Population) | 1.6 | 7.85E-04 | 1.7 | 1.28E-04 | 1.5 | 1.24E-03 |
| | Line(Block) | 7.7 | 5.41E-08 | 7.9 | 2.77E-08 | 1.6 | 1.54E-01 |
| | Column(Block) | 4.3 | 3.06E-04 | 3.8 | 1.03E-03 | 2.9 | 7.89E-03 |
| OTU4_*Com* _1 | Block | 3.0 | 1.13E-02 | 2.8 | 1.74E-02 | 6.2 | 1.18E-05 |
| | Population | 3.7 | 1.15E-15 | 3.6 | 1.16E-14 | 5.0 | 4.34E-24 |
| | Accession(Population) | 1.5 | 1.50E-03 | 1.5 | 1.32E-03 | 2.0 | 6.72E-07 |
| | Line(Block) | 4.1 | 4.93E-04 | 4.4 | 2.13E-04 | 2.1 | 4.85E-02 |
| | Column(Block) | 1.0 | 4.44E-01 | 1.1 | 3.76E-01 | 1.0 | 3.92E-01 |
| OTU5_*Pmo* _1 | Block | 22.0 | 1.72E-20 | 26.2 | 2.75E-24 | 6.2 | 1.36E-05 |
| | Population | 3.9 | 7.07E-17 | 4.0 | 1.57E-17 | 6.0 | 1.53E-30 |
| | Accession(Population) | 1.6 | 8.09E-04 | 1.6 | 2.82E-04 | 2.0 | 1.40E-07 |
| | Line(Block) | 8.9 | 2.34E-09 | 10.7 | 2.05E-11 | 0.6 | 7.18E-01 |
| | Column(Block) | 0.8 | 6.06E-01 | 0.6 | 7.60E-01 | 1.0 | 4.02E-01 |
| OTU5 _*Pmo* _2 | Block | 18.3 | 4.86E-17 | 19.2 | 7.01E-18 | 10.6 | 9.58E-10 |
| | Population | 3.7 | 1.12E-15 | 3.8 | 6.08E-16 | 5.0 | 6.86E-24 |
| | Accession(Population) | 2.0 | 8.49E-07 | 1.8 | 2.42E-05 | 1.8 | 9.27E-06 |
| | Line(Block) | 4.8 | 8.10E-05 | 5.1 | 3.84E-05 | 1.3 | 2.60E-01 |
| | Column(Block) | 1.9 | 7.95E-02 | 1.3 | 2.78E-01 | 3.1 | 5.92E-03 |
| OTU6_*Psi* _1 | Block | 88.1 | 2.68E-71 | 97.4 | 2.72E-77 | 3.7 | 2.54E-03 |
| | Population | 4.2 | 9.50E-19 | 3.7 | 3.55E-15 | 6.0 | 2.17E-30 |
| | Accession(Population) | 1.6 | 7.53E-04 | 1.6 | 2.18E-04 | 1.5 | 2.46E-03 |
| | Line(Block) | 2.6 | 1.57E-02 | 4.1 | 4.21E-04 | 1.9 | 7.49E-02 |
| | Column(Block) | 0.3 | 9.14E-01 | 0.4 | 9.06E-01 | 1.1 | 3.73E-01 |
| OTU6_*Psi* _2 | Block | 16.2 | 4.57E-15 | 24.2 | 2.15E-22 | 5.4 | 7.19E-05 |
| | Population | 3.4 | 8.04E-14 | 3.2 | 3.60E-12 | 4.6 | 3.71E-21 |
| | Accession(Population) | 1.6 | 1.10E-03 | 1.6 | 1.03E-03 | 2.1 | 8.41E-08 |
| | Line(Block) | 2.8 | 1.04E-02 | 4.1 | 4.27E-04 | 2.2 | 4.49E-02 |
| | Column(Block) | 0.6 | 7.67E-01 | 0.7 | 6.49E-01 | 0.5 | 7.72E-01 |
| OTU13_*M* sp_1 | Block | 64.8 | 5.40E-55 | 63.2 | 8.49E-54 | 5.7 | 4.01E-05 |
| | Population | 3.7 | 1.87E-15 | 3.5 | 2.28E-14 | 4.9 | 6.31E-23 |
| | Accession(Population) | 1.3 | 4.09E-02 | 1.2 | 1.03E-01 | 1.8 | 3.24E-05 |
| | Line(Block) | 10.6 | 3.01E-11 | 13.8 | 8.95E-15 | 2.7 | 1.40E-02 |
| | Column(Block) | 0.5 | 7.99E-01 | 0.5 | 7.74E-01 | 0.8 | 5.64E-01 |
| OTU13_*M* sp_2 | Block | 47.9 | 2.33E-42 | 39.4 | 1.68E-35 | 44.8 | 1.69E-39 |
| | Population | 5.1 | 3.96E-25 | 5.3 | 5.96E-26 | 5.9 | 8.06E-30 |
| | Accession(Population) | 1.8 | 2.36E-05 | 1.9 | 1.85E-06 | 2.2 | 2.01E-08 |
| | Line(Block) | 3.9 | 7.86E-04 | 4.3 | 2.93E-04 | 1.6 | 1.55E-01 |
| | Column(Block) | 1.0 | 4.27E-01 | 1.3 | 2.73E-01 | 1.2 | 2.97E-01 |
| OTU29_*Sph* _1 | Block | 35.8 | 1.34E-32 | 43.4 | 7.27E-39 | 6.2 | 1.19E-05 |
| | Population | 5.1 | 1.07E-24 | 4.4 | 1.75E-20 | 5.9 | 1.28E-29 |
| | Accession(Population) | 1.9 | 2.08E-06 | 1.9 | 3.02E-06 | 1.6 | 7.44E-04 |
| | Line(Block) | 4.5 | 1.82E-04 | 4.6 | 1.27E-04 | 3.4 | 2.80E-03 |
| | Column(Block) | 1.9 | 8.66E-02 | 1.6 | 1.32E-01 | 2.5 | 1.93E-02 |
| OTU29_*Sph* _2 | Block | 39.2 | 1.98E-35 | 40.8 | 1.07E-36 | 11.7 | 7.11E-11 |
| | Population | 4.8 | 6.51E-23 | 4.2 | 5.26E-19 | 5.7 | 2.30E-28 |
| | Accession(Population) | 2.0 | 4.74E-07 | 1.9 | 4.34E-06 | 1.8 | 1.66E-05 |
| | Line(Block) | 5.6 | 1.23E-05 | 5.9 | 4.93E-06 | 1.5 | 1.60E-01 |
| | Column(Block) | 0.7 | 6.71E-01 | 0.6 | 7.08E-01 | 0.3 | 9.25E-01 |

**Supplementary Table 2.** Differences among bacterial species (OTUs) and between strains within a given bacterial species for the response of three reproductive traits to the individual inoculation of 13 bacterial strains. Linear models were run with the function *lm* of the library *lme4* under the R environment.

| | | Total seed production | | Fruit number | | Mean fruit length | |
|---|---|---|---|---|---|---|---|
| | Model terms | *F* | *P* | *F* | *P* | *F* | *P* |
| Full model | OTU | 38.1 | 6.43E-44 | 49.1 | 2.08E-56 | 20.2 | 4.44E-23 |
| | strain(OTU) | 16.7 | 7.45E-19 | 21.8 | 6.15E-25 | 3.4 | 2.73E-03 |
| Per OTU | | | | | | | |
| OTU2 | strain | 11.7 | 7.24E-04 | 10.7 | 1.17E-03 | 1.9 | 1.66E-01 |
| OTU3a | strain | 0.0 | 9.38E-01 | 0.0 | 9.89E-01 | 0.6 | 4.35E-01 |
| OTU5 | strain | 4.1 | 4.44E-02 | 9.9 | 1.81E-03 | 11.0 | 1.04E-03 |
| OTU6 | strain | 8.7 | 3.41E-03 | 13.2 | 3.22E-04 | 0.1 | 8.16E-01 |
| OTU13 | strain | 25.9 | 6.36E-07 | 34.0 | 1.40E-08 | 1.0 | 3.29E-01 |
| OTU29 | strain | 92.0 | 3.45E-19 | 114.8 | 6.45E-23 | 6.1 | 1.44E-02 |

**Supplementary Table 3.** Relationship at the accession level between the phenotypic response to the individual inoculation of 13 bacterial strains and phenotypic variation in the absence of inoculation (mock) for the three reproductive traits. Analysis of covariance was run with the function *lm* of the library *lme4* under the R environment. $R^2$: percentage of variance explained by model terms.

| | | Total seed production | | | Fruit number | | | Mean fruit length | | |
|---|---|---|---|---|---|---|---|---|---|---|
| | Model terms | *F* | *P* | $R^2$ | *F* | *P* | $R^2$ | *F* | *P* | $R^2$ |
| Full model | mock | 214.1 | 4.43E-46 | | 177.9 | 6.81E-39 | | 126.3 | 1.97E-28 | |
| | OTU | 43.0 | 1.68E-49 | | 54.4 | 2.71E-62 | | 21.6 | 1.13E-24 | |
| | mock × OTU | 7.9 | 2.04E-08 | | 6.9 | 3.02E-07 | | 1.6 | 1.50E-01 | |
| | strain(OTU) | 18.8 | 2.38E-21 | | 24.1 | 1.14E-27 | | 3.6 | 1.63E-03 | |
| | mock × strain(OTU) | 0.5 | 8.36E-01 | | 0.9 | 5.14E-01 | | 1.5 | 1.80E-01 | |
| Per OTU | | | | | | | | | | |
| OTU2 | mock | 53.3 | 2.60E-12 | | 49.8 | 1.19E-11 | | 33.3 | 1.94E-08 | |
| | strain | 13.5 | 2.83E-04 | | 12.4 | 5.09E-04 | | 2.1 | 1.47E-01 | |
| | mock × strain | 0.6 | 4.27E-01 | | 1.5 | 2.21E-01 | | 4.4 | 3.74E-02 | |
| OTU3a | mock | 10.3 | 1.51E-03 | | 9.3 | 2.52E-03 | | 2.4 | 1.26E-01 | |
| | strain | 0.0 | 9.37E-01 | | 0.0 | 9.89E-01 | | 0.6 | 4.37E-01 | |
| | mock × strain | 0.8 | 3.62E-01 | | 0.2 | 6.44E-01 | | 0.4 | 5.32E-01 | |
| OTU5 | mock | 59.1 | 2.18E-13 | | 48.2 | 2.41E-11 | | 27.6 | 2.79E-07 | |
| | strain | 4.8 | 2.84E-02 | | 11.4 | 8.23E-04 | | 11.8 | 6.71E-04 | |
| | mock × strain | 0.2 | 6.82E-01 | | 0.0 | 8.63E-01 | | 0.0 | 9.06E-01 | |
| OTU6 | mock | 47.4 | 3.33E-11 | | 39.0 | 1.48E-09 | | 33.0 | 2.27E-08 | |
| | strain | 10.0 | 1.69E-03 | | 15.0 | 1.34E-04 | | 0.1 | 7.96E-01 | |
| | mock × strain | 0.6 | 4.57E-01 | | 2.6 | 1.11E-01 | | 0.2 | 6.71E-01 | |
| OTU13 | mock | 18.1 | 2.84E-05 | | 14.6 | 1.63E-04 | | 14.7 | 1.56E-04 | |
| | strain | 27.3 | 3.26E-07 | | 35.5 | 7.23E-09 | | 1.0 | 3.14E-01 | |
| | mock × strain | 0.1 | 7.28E-01 | | 0.0 | 8.79E-01 | | 0.2 | 6.96E-01 | |
| OTU29 | mock | 7.1 | 8.01E-03 | | 3.8 | 5.34E-02 | | 22.4 | 3.45E-06 | |
| | strain | 93.6 | 1.88E-19 | | 115.5 | 5.30E-23 | | 6.6 | 1.08E-02 | |
| | mock × strain | 0.3 | 5.82E-01 | | 0.0 | 9.79E-01 | | 5.0 | 2.62E-02 | |
| Per strain | | | | | | | | | | |
| OTU2_*Pfu* _1 | mock | 25.9 | 1.07E-06 | 14.7 | 26.6 | 7.72E-07 | 15.1 | 6.9 | 9.71E-03 | 4.4 |
| OTU2_*Pfu* _2 | mock | 28.9 | 2.85E-07 | 16.3 | 23.9 | 2.55E-06 | 13.8 | 30.7 | 1.36E-07 | 17.2 |
| OTU3a_*Oxa* _1 | mock | 1.9 | 1.66E-01 | 1.3 | 2.6 | 1.11E-01 | 1.7 | 0.3 | 5.83E-01 | 0.2 |
| OTU3a_*Oxa* _2 | mock | 13.0 | 4.16E-04 | 8.0 | 8.8 | 3.46E-03 | 5.6 | 3.6 | 5.85E-02 | 2.4 |
| OTU4_*Com* _1 | mock | 33.9 | 3.47E-08 | 18.4 | 30.8 | 1.25E-07 | 17.1 | 13.2 | 3.90E-04 | 8.1 |
| OTU5_*Pmo* _1 | mock | 43.0 | 8.38E-10 | 22.3 | 39.6 | 3.21E-09 | 20.9 | 16.8 | 6.74E-05 | 10.1 |
| OTU5_*Pmo* _2 | mock | 23.7 | 2.85E-06 | 13.6 | 17.8 | 4.24E-05 | 10.6 | 11.9 | 7.40E-04 | 7.4 |
| OTU6_*Psi* _1 | mock | 22.4 | 4.99E-06 | 13.0 | 12.7 | 4.96E-04 | 7.8 | 18.8 | 2.67E-05 | 11.2 |
| OTU6_*Psi* _2 | mock | 25.1 | 1.49E-06 | 14.4 | 26.7 | 7.38E-07 | 15.1 | 14.3 | 2.20E-04 | 8.7 |
| OTU13_*M* sp_1 | mock | 11.0 | 1.13E-03 | 6.8 | 8.3 | 4.49E-03 | 5.3 | 5.1 | 2.59E-02 | 3.3 |
| OTU13_*M* sp_2 | mock | 7.3 | 7.60E-03 | 4.7 | 6.4 | 1.25E-02 | 4.1 | 10.7 | 1.32E-03 | 6.7 |
| OTU29_*Sph* _1 | mock | 3.7 | 5.54E-02 | 2.4 | 1.3 | 2.54E-01 | 0.9 | 26.0 | 9.95E-07 | 14.8 |
| OTU29_*Sph* _2 | mock | 3.7 | 5.67E-02 | 2.4 | 3.2 | 7.57E-02 | 2.1 | 2.9 | 8.99E-02 | 1.9 |

**Supplementary Table 4.** Fold enrichment (FE) of the top SNPs in the 0.5% upper tail of the genome-wide spatial differentiation (XtX) distribution for each reproductive trait in response to the individual inoculation of 13 bacterial strains.

| | Total seed production | | | Fruit number | | | Mean fruit length | | |
|---|---|---|---|---|---|---|---|---|---|
| Strain | No. Top SNPs | FE | *P* | No. Top SNPs | FE | *P* | No. Top SNPs | FE | *P* |
| OTU2_*Pfu* _1 | 15 | 2.04 | 0.0035 | 23 | 4.24 | 0.0000 | 2 | 0.66 | 0.1830 |
| OTU2_*Pfu* _2 | 6 | 1.55 | 0.0445 | 8 | 1.48 | 0.0317 | 22 | 5.45 | 0.0000 |
| OTU3a_*Oxa* _1 | 3 | 1.55 | 0.0536 | 4 | 1.90 | 0.0263 | 2 | 0.73 | 0.2005 |
| OTU3a_*Oxa* _2 | 28 | 3.45 | 0.0000 | 24 | 3.73 | 0.0000 | 8 | 1.05 | 0.0220 |
| OTU4_*Com* _1 | 3 | 1.19 | 0.0843 | 4 | 1.28 | 0.0526 | 11 | 1.67 | 0.0177 |
| OTU5_*Pmo* _1 | 14 | 1.74 | 0.0214 | 22 | 2.80 | 0.0000 | 5 | 0.75 | 0.1329 |
| OTU5_*Pmo* _2 | 0 | 0.00 | 0.6873 | 13 | 1.99 | 0.0096 | 9 | 1.34 | 0.0289 |
| OTU6_*Psi* _1 | 1 | 0.69 | 0.1839 | 8 | 4.23 | 0.0012 | 0 | 0.00 | 0.4065 |
| OTU6_*Psi* _2 | 6 | 2.84 | 0.0032 | 10 | 2.05 | 0.0023 | 4 | 1.26 | 0.0301 |
| OTU13_*M* sp_1 | 12 | 1.56 | 0.0098 | 11 | 2.56 | 0.0049 | 8 | 2.74 | 0.0015 |
| OTU13_*M* sp_2 | 1 | 0.85 | 0.1763 | 29 | 4.63 | 0.0000 | 15 | 1.48 | 0.0038 |
| OTU29_*Sph* _1 | 4 | 0.62 | 0.1081 | 5 | 0.93 | 0.0540 | 7 | 2.10 | 0.0226 |
| OTU29_*Sph* _2 | 7 | 2.69 | 0.0044 | 5 | 1.80 | 0.0385 | 3 | 1.09 | 0.0621 |

**Supplementary Table 5.** Enrichment for the number of candidate genes in the 5% upper and lower tail of the Tajima's D distribution for each reproductive trait in response to the individual inoculation of 13 bacterial strains. In agreement with the composite LR test (Dataset S7), the 5% lower tail of Tajima's D distribution (also suggestive of selective sweeps) displayed no significant enrichment for candidate genes identified for each 'reproductive trait × bacterial strain' combination (SI Appendix Table S5).

| | | Total seed production | | Fruit number | | Mean fruit length | |
|---|---|---|---|---|---|---|---|
| Tajima's D distribution | Strain | No. Genes | *P* | No. Genes | *P* | No. Genes | *P* |
| 5% upper tail | OTU2_*Pfu* _1 | 23 | 0.0000 | 14 | 0.0003 | 3 | 0.2653 |
| | OTU2_*Pfu* _2 | 7 | 0.0949 | 8 | 0.1211 | 8 | 0.0135 |
| | OTU3a_*Oxa* _1 | 5 | 0.0283 | 2 | 0.5634 | 9 | 0.0108 |
| | OTU3a_*Oxa* _2 | 14 | 0.0193 | 14 | 0.0076 | 12 | 0.0044 |
| | OTU4_*Com* _1 | 5 | 0.1296 | 4 | 0.2786 | 9 | 0.1491 |
| | OTU5_*Pmo* _1 | 12 | 0.0305 | 4 | 0.6022 | 12 | 0.0041 |
| | OTU5_*Pmo* _2 | 10 | 0.0029 | 15 | 0.0034 | 6 | 0.1917 |
| | OTU6_*Psi* _1 | 1 | 0.8714 | 1 | 0.8862 | 4 | 0.1193 |
| | OTU6_*Psi* _2 | 2 | 0.5506 | 3 | 0.5465 | 5 | 0.0744 |
| | OTU13_*M* sp_1 | 12 | 0.0303 | 8 | 0.0497 | 7 | 0.0345 |
| | OTU13_*M* sp_2 | 4 | 0.1312 | 11 | 0.0440 | 26 | 0.0000 |
| | OTU29_*Sph* _1 | 4 | 0.5306 | 4 | 0.5274 | 4 | 0.4275 |
| | OTU29_*Sph* _2 | 3 | 0.3181 | 8 | 0.0265 | 4 | 0.0342 |
| 5% lower tail | OTU2_*Pfu* _1 | 0 | 1.0000 | 0 | 1.0000 | 0 | 1.0000 |
| | OTU2_*Pfu* _2 | 4 | 0.4890 | 4 | 0.6606 | 1 | 0.9191 |
| | OTU3a_*Oxa* _1 | 1 | 0.7559 | 1 | 0.8371 | 0 | 1.0000 |
| | OTU3a_*Oxa* _2 | 3 | 0.9526 | 0 | 1.0000 | 1 | 0.9834 |
| | OTU4_*Com* _1 | 0 | 1.0000 | 0 | 1.0000 | 5 | 0.6662 |
| | OTU5_*Pmo* _1 | 5 | 0.6405 | 3 | 0.7737 | 2 | 0.8997 |
| | OTU5_*Pmo* _2 | 1 | 0.9337 | 2 | 0.9693 | 4 | 0.4948 |
| | OTU6_*Psi* _1 | 2 | 0.6411 | 2 | 0.6822 | 1 | 0.7920 |
| | OTU6_*Psi* _2 | 0 | 1.0000 | 1 | 0.9313 | 1 | 0.8558 |
| | OTU13_*M* sp_1 | 5 | 0.6609 | 2 | 0.8625 | 7 | 0.0309 |
| | OTU13_*M* sp_2 | 3 | 0.2707 | 2 | 0.9480 | 1 | 0.9934 |
| | OTU29_*Sph* _1 | 1 | 0.9703 | 1 | 0.9697 | 1 | 0.9548 |
| | OTU29_*Sph* _2 | 0 | 1.0000 | 0 | 1.0000 | 0 | 1.0000 |

**Supplementary Table 6.** List of the seven candidate genes for total seed production with signatures of local adaptation, signatures of balancing selection, and belonging to enriched biological processes.

| ATG number | Annotation | Subcategory | Function | References | Seed germination* | Flower stage* | Leaf* | Seeds at flower Abscission* | Pods / Seeds at silique stage* |
|---|---|---|---|---|---|---|---|---|---|
| *AT1G32870* | ANAC013 | NAC transcription factor | Role in abiotic stress signaling linked to oxidative stress and seed germination | (Jurdak et al. 2021; Eysholdt-Derzsó et al. 2023) | X | | | | X |
| *AT1G51840* | SIF1 | Kinase-like protein (LRR-RLK), stress-induced factor | Role in abiotic stress signaling linked to drought and salt osmotic stress | (Yuan et al. 2018) | X | | | | |
| *AT2G36910* | ABCB1 | ATP-binding cassette (ABC) transporters | Role in auxin transport in the stamen | (Liu et al. 2022) | | X | | X | |
| *AT3G05480* | RAD9 | Regulation of DNA repair | Involved in the regulation of DNA damage repair and homologous recombination | (Heitzeberg et al. 2004) | | X | | | |
| *AT5G01550* | LecRKA4.2 | Lectin receptor kinase subfamily A4 | Role in the negative regulation of abscisic acid response in seed germination | (Xin et al. 2009) | | X | X | | |
| *AT5G01560* | LecRKA4.3 | Lectin receptor kinase subfamily A4 | Role in the negative regulation of abscisic acid response in seed germination | (Xin et al. 2009) | | | X | X | X |
| *AT5G49490* | AGL83 | AGAMOUS-like MADS box transcription factor | Weakly expressed in the female gametophyte | (Bemer et al. 2010) | | X | | | |

***** Information extracted from TAIR (https://www.arabidopsis.org/), BAR eFP Browser: localization and presentation of the developmental stages at which these genes are most highly expressed.
